## Supplemental Figure 1 for "Advanced Biophysical Model to Capture Channel Variability for EQS Capacitive HBC"

Arunashish Datta, Mayukh Nath, David Yang and Shreyas Sen, *Senior Member, IEEE*

School of Electrical and Computer Engineering, Purdue University

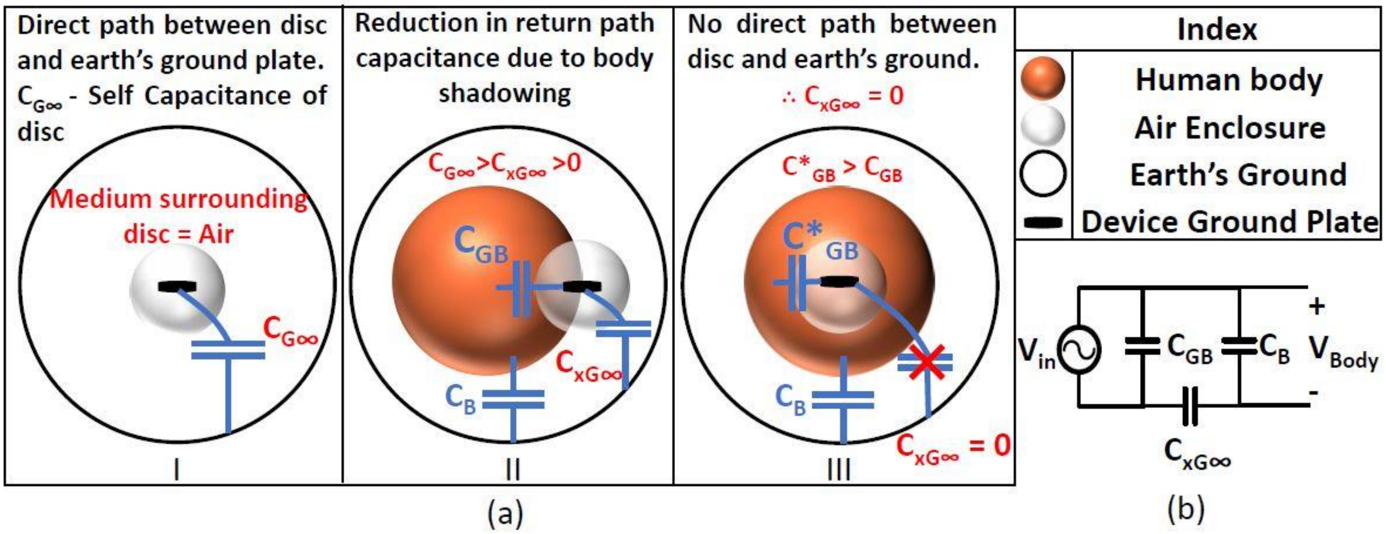


Fig. S1. (a) Sphere model developed to understand the effects of body to floating ground plate capacitance ($C_{GB}$) on the return path capacitance ($C_{xG\infty}$). As shown in the index, the first sphere depicts a simplified human body structure. The next sphere represents an air enclosure whereas the outer hollow sphere represents the earth's ground plane. A disc is used to represent the floating ground plate of a device. In figure I, the medium between the outer sphere representing earth's ground in air. Hence, the disc present in an air enclosure has a direct path to the earth's ground and the capacitance between them will be equal to the self capacitance of a disc ($C_{G\infty=8\epsilon_0 a}$) where the radius of the disc ($a$) is much lesser than the radius of the outer sphere. Figure II represents a situation where the air enclosure and the disc are partially inside the human body. We still have a direct path between the disc and the earth's ground. However, the disc is also shadowed by the body resulting in the body to floating ground plate capacitance ($C_{GB}$) which reduces the return path capacitance ($C_{xG\infty}$). $C_B$ represents the capacitance between body and earth's ground. Figure III represents the case where the air enclosure with the disc is completely inside the body. In this case, there is no direct path between the disc and the earth's ground which reduces the return path capacitance to zero. (b) A simplified circuit model is developed for a capacitive HBC channel with respect to the amount of signal being coupled to the body.

### I. SUPPLEMENTAL MATERIAL

The effect of fringe fields due to body shadowing is modeled by the body to floating ground plate capacitance ($C_{GB}$) in the advanced biophysical model developed. However, the capacitance $C_{GB}$ is present parallel to the transmitter ($V_{in}$) in the circuit model as shown in Fig. S1 (b). Hence, it doesn't directly affect the value of the potential received at the body as shown by Eqn: S1.

$$\frac{V_{Body}}{V_{in}} \approx \frac{C_{xG\infty}}{C_B + C_{xG\infty}} \tag{S1}$$

However, the presence of body to floating ground plate capacitance has an effect on the return path capacitance ($C_{G\infty}$) of the system. Due to presence of fringe fields between the floating ground plate of the transmitter and the body, the number of field lines between from the floating ground plate of the transmitter and the earth's ground plane reduces. Thus, the fringe fields effectively steal some field lines from the floating ground plate to the earth's ground causing the return path capacitance to decrease. This can be better understood using the sphere thought model shown in Fig. S1 (a).

This simplified model is used to draw an analogy to the capacitive HBC system. The human body is modelled by the red sphere. A smaller sphere which is an air enclosure surrounds the disc which represents the floating ground plate of the transmitter or receiver device. The medium outside the spheres also consists of air which is further surrounded by the earth's ground represented by the large circle.

In Fig. S1 (a) I, the air enclosure with the disc is kept far apart from the human body (red sphere). In this situation, the disc has a direct path to the earth's ground outside as the medium present between them is air. In this scenario, the only capacitance we observe is present between the disc and the earth's ground which is the self capacitance of the disc

$(C_{G\infty} = 8\epsilon_0 a)$ when the radius of the disc $(a)$ is much lesser than the radius of the larger sphere representing earth's ground. This is a corner case where the human body and the floating ground plate of the device are placed far away from each other. This ensures that the human body does not shadow the device and the return path capacitance is at a maximum value.

Fig. S1 (a) III represents another corner case where the disc present within the air enclosure is completely surrounded by the red sphere representing the body. In this case, the disc can be visualized to be an implantable device present within the body. Here, a capacitance between the disc and the body $(C*_{GB})$ is observed. A capacitance between the body and the earth's ground $(C_B)$ is also present as observed in a typical capacitive HBC scenario. However, since the smaller sphere (air enclosure) is completely surrounded by the body, there are no field lines escaping from the disc directly to the earth's ground plane. Hence, the component $C_{G\infty}$ seen in Fig. S1 (a) I has reduced to $0$ in Fig. S1 (a) III.

Fig. S1 (a) II falls between the two corner cases shown previously. This replicates the instance for a wearable device in capacitive HBC. Here, the disc and the air enclosure are not completely surrounded by the body. We again observe a capacitance between the body and earth's ground plate $(C_B)$. Further, we see that there is a capacitance $(C_{GB})$ between the disc and the body for the part of the sphere where the air enclosure is in contact with the body. The body to disc capacitance present in this case is less than the capacitance between the disc and body for Fig. S1 (a) III $(C*_{GB})$ as the surface area of body capacitively coupling to the disc is lesser in Fig. S1 (a) II. Lastly, since there is now a direct path for the field lines to travel from the disc to the environment outside the sphere system, we have a capacitance $(C_{xG\infty})$ from the disc to infinity. This capacitance is lesser than the self capacitance of a disc observed in Fig. S1 (a) I as the surface area of the disc exposed to the earth's ground is lesser in the second case. This phenomenon of the return path capacitance reduction due to the disc being in close proximity to the body is termed as the body shadowing effect. The sphere model illustrates that the return path capacitance is not constant at all positions of the human body and is also lesser than the self capacitance of the disc whenever the ground plate is close to the human body.

### II. Key Takeaway

A thought model is shown where the effect of body shadowing is explained. The effect of body to floating ground plate fringe capacitance on the reduction of return path capacitance is explained. The increase in fringe capacitance results in decreasing return path capacitance. This effect of fringe fields stealing field lines away from the direct path between floating ground plate of a device (transmitter or receiver) and the earth's ground plate is called the body's shadowing effects.
